# Parental effects via glyphosate-based herbicides in a bird model?

**DOI:** 10.1101/2019.12.21.885855

**Authors:** Suvi Ruuskanen, Miia Rainio, Maiju Uusitalo, Kari Saikkonen, Marjo Helander

**Affiliations:** Department of Biology, University of Turku, Vesilinnantie 5, 20500, Turku, Finland; Biodiversity Unit, University of Turku, Vesilinnantie 5, 20500, Turku, Finland

**Keywords:** herbicide, glyphosate, maternal effect, oxidative status

## Abstract

Controversial glyphosate-based herbicides (GBHs) are the most frequently used herbicides across the globe. In an increasing number of studies, researchers have identified GBH residues in soil, water, crops, and food products exposing non-target organisms to health risks; these organisms include wildlife, livestock, and humans. However, GBH-related parental effects are poorly understood. In the case of birds, GBHs may be transferred directly from mother to developing offspring (i.e. direct effects) via eggs, or they may indirectly influence offspring performance by altered maternal condition or resource allocation to eggs, for example. We experimentally exposed a parental generation of Japanese quails (*Coturnix japonica*) to GBHs or respective controls, recorded egg quality and glyphosate residues in eggs, and studied embryonic development and oxidative biomarkers. Glyphosate accumulated in eggs (ca 0.76 kg/mg). Embryonic development tended to be lower in eggs of GBH-exposed parents compared to control parents. Embryonic brain tissue from GBH-exposed parents tended to express more lipid damage. Given that we detected no differences in egg quality (egg, yolk, or shell mass, or egg hormone concentration) across the treatment groups, our results suggest these are likely direct effects of GBHs on offspring rather than indirect effects via altered maternal allocation of resources or hormonal signals.

**Capsule:** Experimental, long-term parental exposure to GBHs tends to hinder offspring embryonic development and increase embryonic oxidative damage to lipids in a bird model.

## Introduction

Glyphosate (N-[Phosphonomethyl]glycine)-based herbicides (GBHs) are the most frequently used herbicides globally and one of the most controversial agrochemicals (Benbrook, 2016). Evidence is accumulating with regard to the potentially negative effects of glyphosate on the development, phenotype, and fitness of virtually all non-target animal taxa from invertebrates to vertebrates (Gill et al., 2018; Szekacs & Darvas, 2018; Van Bruggen et al., 2018). Non-target organisms are commonly exposed to GBH residues in the food chain because residues can persist in soil, water, and plants (Bai & Ogbourne, 2016; Helander et al., 2012). Consequently, different regulatory authorities heatedly debate the effects of GBH in our ecosystems.

Organisms in early developmental stages are generally more susceptible to external stress compared to adults. In the case of environmental toxins, this may be related to disturbed ontogeny or undeveloped detoxification metabolism in juveniles. In aquatic animals, embryos can be directly exposed to GBHs via the surrounding water. Glyphosate and commercial products (e.g. Roundup®) made with glyphosate have been repeatedly reported to cause embryo mortality and deformations in fish (summarized e.g. in Burella et al., 2018; Schweizer et al., 2019; Webster et al., 2014) and aquatic amphibians (Babalola et al., 2019; Paganelli, et al., 2010). In contrast, mammal and bird embryos and fetuses are exposed to glyphosate residues only via maternal transfer of the chemicals, which results in malformations, altered sex ratios, and low sperm quality in rodent models (Dallegrave et al., 2003; Dallegrave et al., 2007; Ren et al., 2018). Such effects are referred as (transmissive) maternal effects (Marshall & Uller, 2007). Furthermore, recent studies suggest that effects of GBHs on the next generation can be mediated via epigenetic paternal effects, for example via alternations of paternal sperm (Kubsad et al., 2019; Mesnage et al., 2015).

However, the true maternal and paternal effects of GBH are poorly understood because the majority of the studies are those involving direct embryo manipulations with high doses of GBHs. The authors of future studies should take into account that GBHs may influence the quality of the resources allocated to eggs/embryos and thus offspring development, phenotype, and fitness *indirectly*. Prenatal environmental conditions, and for example hormonal signals from the mother are known to have crucial importance for offspring development and even lasting effects into adulthood (Ruuskanen, 2015; Ruuskanen and Hsu, 2018; Moore et al., 2019, Yin et al., 2019).

In this study we used birds as a model to study the parental and developmental effects of GBHs. Birds are highly underrepresented in studies testing the adverse effects of GBH residues on non-target taxa (Gill et al., 2018), although they have recently been suggested as a key group for biomonitoring with regard to the effects of GBHs (Kissane & Shephard, 2017). The importance of poultry in food production also calls for more attention on the effects of GBHs in birds. In the two available studies of poultry and GBH-related maternal effects, a direct injection of a relatively high concentration of Roundup (10 mg/kg glyphosate) was found to decrease hatchability, induce oxidative stress and cause damage to lipids in the exposed chicks, as compared to the control group (Fathi et al., 2019), potentially via the disruption of retinoid acid signaling (Paganelli et al., 2010).

To understand the potential for GBH-induced parental effects, we studied parental exposure of GBHs on embryo development and key physiological biomarkers—embryonic brain oxidative status in a bird model. To our knowledge, this is the first long-term study on parental effects of GBHs in bird taxa. Oxidative stress refers to the imbalance between reactive oxygen species (ROS) and antioxidants: If antioxidants are not able to neutralize ROS, oxidative damage to cell components (proteins, lipids, and DNA) will occur, which then has negative consequences on cell functions (Halliwell & Gutteridge, 2015). GBHs have been previously found to induce oxidative stress and damage in a variety of organisms and tissues, including embryos (reviewed in Gill et al., 2018). We quantified glyphosate residues in eggs, but also maternal allocation to eggs (egg, yolk, and shell mass and yolk thyroid hormone concentration) to account for potential indirect GBH effects. Prenatal thyroid hormones (THs) (thyroxine, T4 and triiodothyronine, T3) play a key role in coordinating embryo development (e.g. Ruuskanen and Hsu, 2018; Ruuskanen et al., 2016), especially brain development (Darras et al., 2009). Embryo THs have been reported to vary with maternal GBH exposure in rats (de Souza et al., 2017), but generally the effects of GBHs on THs are poorly understood. Japanese quails were selected as the model species because the results can be applied to both wild birds feeding on GBH-contaminated food in the field and to poultry farming. We experimentally exposed parental bird generation to GBHs or respective controls from 10 days of age to 12 months. The egg samples were collected at 4 and 12 months to examine the potential cumulative effects of long-term exposure. We predicted (1) negative effects on egg quality (egg, yolk, and shell mass as well as egg thyroid hormones); (2) negative effects on embryo development; and (3) higher oxidative stress and damage in the embryos from GBH-exposed parents compared to controls.

## Material and methods

We performed an experiment in which a parental generation of Japanese quails were fed with either GBH-contaminated food (N = 13 breeding pairs) or control food (N = 13 breeding pairs) from the age of 10 days to 12 months. Eggs from the parental generation were artificially incubated and analyzed for embryo traits.

### GBH treatments to the parental generation

Details of the experimental design and parental exposures are described in the work of Ruuskanen et al. (2019). The grandparental birds originated from local breeders in Finland. The chicks of the parental generation were randomly and evenly divided into two groups from all parents and sexes. Due to unknown reasons, the sex ratio of hatchlings was biased toward males; however, all hatched chicks were included in the experiment. The samples sizes were: 15 (GBH, female), 23 (GBH, male), 14 (control, female), and 24 (control, male), from which 13 breeding pairs for each treatment were formed.

The GBH-exposed group was fed organic food (Organic food for laying poultry, “Luonnon Punaheltta” Danish Agro, Denmark) with added commercial GBH (Roundup Flex^®^ 480g/l glyphosate, present as 588g/l [43.8% w.w] of potassium salt of glyphosate, with surfactants alkylpolyglycoside (5% of weight) and nitrotryl (1% of weight) (AXGD42311 5/7/2017, Monsanto, 2002). The control group was fed the same organic food in which water was added without GBH. A GBH product was selected over pure glyphosate to mimic the exposures in natural environments including exposure to adjuvants, as adjuvants may increase the toxicity of glyphosate (Gill et al., 2018; Mesnage & Antoniou, 2018). However, with this experimental design we could not distinguish the potential effects of adjuvants themselves, or whether they altered the effects of the active ingredient, glyphosate. The concentration of glyphosate in the GBH food was aimed at ca 200 mg/kg food, which is ca ½ of that calculated in grains (available to granivorous birds) after GBHs are spread on a grain field (Eason & Scanlon, 2002). This concentration in the food corresponds to a dose of 12-20 mg glyphosate/kg body mass/day in full-grown Japanese quails. The European Food Safety Authority (EFSA) reports a NOAEL (No Adverse Effects Level) of 100 mg/kg body mass/day for poultry (EFSA, 2018); therefore, our experiment tests a rather moderate concentration well below this threshold. Furthermore, a dose of 347 mg/kg did not negatively influence adult body mass in Japanese quails in a short-term experiment (Eason & Scanlon, 2002). According to the manufacturer, acute toxicity (LC50) (via food) of Roundup Flex® is >4640 mg/kg food for mallards (*Anas platyrhynchos*) and bobwhite quail (*Colinus virginianus)*.

To verify the treatment levels, glyphosate concentration was measured in 6 batches of food and residue levels were measured in excreta (feces and urine) samples after 12 months of exposure. Excreta of 4 to 6 randomly chosen individuals per treatment and sex were pooled. We analyzed glyphosate residues from 2 control pools (one female and one male pool) and 3 glyphosate pools (1 female and 2 male pools). Glyphosate residues were measured via LC-MS/MS (certified laboratory, Groen Agro Control, Delft, The Netherlands). The extraction was performed with a mixture of water and acidified methanol. The analyses were performed with a liquid chromatography coupled to a tandem quadruple mass spectrometer (LC-MS/MS). The separation was performed with a mixed mode column using a gradient based on a mixture of water and acetonitrile. A specific MRM (multiple reaction monitoring) was used to identify the component and standard addition to quantify the concentration. The detection limit was 0.01 mg/kg. The average glyphosate concentration of 6 batches of food was 164 mg/kg (S.E. ± 55 mg/kg). The average glyphosate concentration in 3 pools of excreta samples (urine and fecal matter combined) was 199 mg/kg (S.E. ± 10.5 mg/kg). The control feed and control pools of excreta were free of glyphosate residues (<0.01 mg/kg).

GBH food was prepared every week to avoid potential changes in concentration caused by degradation. Diluted Roundup Flex^®^ was mixed with the organic food in a cement mill (Euro-Mix 125, Lescha, Germany). The food was air-dried and further crushed with a food crusher (Model ETM, Vercella Giuseppe, Italy) to a grain size suitable for the birds considering their age. The control food was prepared using a similar method, but only water was added to the food and a separate cement mill was used (ABM P135 L, Lescha, Germany). After crushing, the dry food was stored in closed containers at 20° C in dry conditions. Separate pieces of equipment for food preparation and storage were used for GBH and control food to avoid contamination.

### Egg collection: Egg mass and egg thyroid hormones

Parental generation was reared in same-sex groups for the first 12 weeks and thereafter in randomly allocated female-male pairs of the same treatment. Eggs were collected when the birds were 4 and 12 months old. Eggs from each cage were collected eggs daily (quails generally lay one egg per day), marked individually, and weighed. A total of 221 and 96 eggs were collected at 4 and 12 months, respectively.

### Egg quality: Yolk and shell mass and thyroid hormones

Yolk and shell mass and yolk thyroid hormones were measured from 12 eggs in the GBH treatment group and 12 eggs in control group, collected after 4 months of exposure. Eggs were thawed and the yolk and shell were separated and weighed (accuracy 1 mg). Yolk was homogenized in MilliQ water (1:1) and a small sample (ca 10 mg) was used for further analysis. Samples were analyzed at the University of Turku for T3 and T4. LCMS/MS was conducted at the facilities of the Turku Center for Biotechnology. T3and T4 were extracted from yolk and plasma following previously published methods (Ruuskanen et al., 2018). T3 and T4 were quantified using a nanoflow liquid chromatography-mass spectrometry (nano-LC-MS/MS) method, developed and validated in the work of Ruuskanen et al. (2018). On-column quantification limits were 10.6 amol for T4 and 17.9 amol for T3. MS data were acquired automatically using Thermo Xcalibur software (Thermo Fisher Scientific) and analyzed using Skyline (MacLean et al., 2010). For the analyses, peak area ratios of sample to internal standard were calculated. TH concentrations, corrected for extraction efficiency, were expressed as pg/mg yolk.

### Glyphosate residue analysis

Details of egg sampling and analysis are presented in detail in the work of Ruuskanen et al. (2019). Fresh eggs collected at 10 months of exposure were frozen at −20° C for glyphosate residue analysis. Prior to analysis, the eggs were thawed and the shells were carefully removed. To avoid contamination, all eggs were processed in a lab that had never been in contact with glyphosate, using clean materials (gloves, petri dishes, and tubes) for each egg. When removing content from each eggshell, the egg content was never in touch with the outer eggshell. Contents of 5 eggs (5 different females) from the control treatment were pooled for glyphosate residue analysis. Contents of 5 eggs (5 different females) from the GBH treatment were individually analyzed for glyphosate residues.

### Egg collection: Development and tissue sampling

Embryo development was assessed after 4 and 12 months of parental exposure to GBHs. At 4 months, 108 GBH and 112 control eggs were collected and artificially incubated for 3 days at 36.8° C and 55% humidity (Rcom Maru Max, Standard CT-190, Autoelex CO. LTD, South-Korea). Eggs were thereafter chilled and assessed for the presence of a normally developed embryo (coded 1) or no embryo/a very small embryo (coded 0).

After parental exposure for 10 months, eggs were collected and artificially incubated for 10 days (i.e. 55% of the normal embryonic developmental period, 17 to 18 days) to detect major developmental defects via histology. A total of 57 embryos were collected, chilled, and washed with 0.9% NaCl and fixed in formalin at +8° C for 16 to 19 days. Thereafter they were dehydrated and fixed in paraffin overnight. Tissue samples were hematoxylin-eosin stained according to standard protocols. Three high quality, randomly selected embryos were photographed using NIS-Elements AR 5.02.00 software. Examples of embryo photographs from parental GBH and control treatments are shown as Supplemental Figures 1a and 1b.

After parental exposure for 12 months, 44 GBH and 52 control eggs were collected fresh and incubated for 10 days as above to assess general development. Thereafter, the brain tissue of the embryo was studied for oxidative status assessment. Embryos were chilled, whole brain tissues were collected, then they were snapped frozen in liquid nitrogen and later stored at −80° C.

The experiments were conducted under licenses from the Animal Experiment Board of the Administrative Agency of South Finland (ESAVI/7225/04.10.07/2017).

### Brain mass and oxidative status biomarkers

We aimed to analyze 2 randomly selected embryo samples per breeding pair (N = 13 pairs/treatment). However, as not all females were producing eggs, or did not produce eggs with (viable) embryos, the final sample size was 19 control embryos (from 10 females) and 16 GBH embryos (from 10 females). Brain homogenates were used to measure oxidative status biomarkers, antioxidant enzymes glutathione-S-transferase (GST), glutathione peroxidases (GPx), catalase (CAT), and oxidative damage to lipids (malonaldehyde, MDA as a proxy, using TBARS assay). Whole brains were weighed (~0.1mg) and homogenized (TissueLyser, Qiagen, Austin, USA) with 200-400 µl KF buffer (0.1 M K_2_HPO_4_ + 0.15 M KCl, pH 7.4). All biomarkers were measured in triplicate (intra-assay coefficient of variability [CV] < 15% in all cases) using an EnVision microplate reader (PerkinElmer, Finland) and calibrated to the protein concentration in the sample. The protein concentration (mg/ml) was measured with a bicinchoninic acid (BCA) protein assay (Smith et al., 1985) using bovine serum albumin (BSA) as a standard (Sigma Chemicals, USA) with the EnVision microplate reader at an absorbance of 570 nm. The GPx-assay (Sigma CGP1) was adjusted from a cuvette to a 384-well plate. GPx was measured following kit instructions but instead of t-Bu-OOH, we used 2 mM H_2_O_2_, which is a substrate for GPx and CAT. To block CAT, 1 mM NaN_3_ was added and the pH was adjusted to 7.0 with the HCl in the buffer provided with the kit (Deisseroth & Dounce, 1970). The change in absorbance was measured at 340 nm. GST-assay (Sigma CS0410) was likewise adjusted from a 96- to a 384-well plate using our own reagents: Dulbecco’s Phosphate Buffered Saline–buffer (DPBS), 200 mM GSH (Sigma G4251), and 100 mM 1-Chloro-2,4-dinitrobenzene (CDNB) (Sigma C6396) in ethanol. The more detailed assay description can be found in the work of Habig et al. (1974). The change in absorbance was measured at 340 nm. The CAT-assay (Sigma CAT100) was adjusted from a cuvette to a 96-well plate. We used a 0.3 mg/ml sample dilution and made our own reagents: 10 × CAT assay buffer (500 mM KF, pH 7.0), CAT dilution buffer (50 mM KF + 0.1% TritonX, pH 7.0), chromogen reagent (0.25 mM 4-aminoantipyrene + 2 mM 3,5-dicloro-2-hydroxybenzenesulfonic acid in 150 mM potassium phosphate buffer, pH 7.0), peroxidase solutions (from horseradish), stop solution (15 mM NaN_3_, Sigma), and 200 mM and 10 mM H_2_O_2_ according to information provided in the technical bulletin (Deisseroth & Dounce, 1970; Fossati et al., 1980). The change in absorbance was measured at 520 nm. The lipid peroxidation was analyzed using a 384-plate modification of a TBARS-assay as described by Espin et al. (2018). In brief, 50 μl of samples diluted in 0.9% NaCl were mixed with 100 μl of TBARS-BHT reagent (15% Trichloroacetic acid, TCA; 0.375% 2-Thiobarbituric acid, TBA and 0.25 N hydrochloric acid, HCl + 2% Butylated hydroxytoluene, BHT) and incubated in a thermoblock at 90° C for 30 min. Samples were then cooled in ice water for 10 min to stop the reaction; they were then centrifuged for 15 min at 2100 g in +6º C. The standard (7 standard points) was prepared from MDA (Sigma Chemicals, USA). The samples were analyzed in black 384-well plates and fluorescence intensity was measured at an excitation/emission wavelength of 530/550 nm.

### Statistical analysis

Egg mass was analyzed using linear mixed models with treatment (GBH or control), exposure duration (4 or 12 months) and their interaction as predictors, female mass as a covariate, and breeding pair ID as a random effect to control for non-independence of eggs from the same pair. Differences between the GBH and control groups in yolk and eggshell mass and egg thyroid hormone levels were analyzed with two-sample t-tests. The likelihood of embryo development was analyzed using generalized linear mixed models (binomial distribution, logit link) with similar predictors. Embryo brain mass, GST, GP, CAT, and MDA were analyzed using linear mixed models with treatment as a predictor and female ID and assay ID (if applicable) as random effects. The Kenward-Rogers method was used to estimate the degrees of freedom. Residuals of the models were visually inspected to confirm normality and heteroscedasticity. All statistical analyses were conducted with SAS Enterprise Guide 7.1. All data are available as Supplementary datafiles (1-3).

## Results

We detected 0.76 mg/kg (S.D. ± 0.16) of glyphosate residue in eggs (see also Ruuskanen et al., 2019), which is above the levels reported in the previous literature (FAO, 2005). Egg mass from GBH and control parents did not differ after 4 or 12 months of exposure (treatment F_1, 17,1_ = 0.12, p = 0.73, treatment*period F_1, 270_ = 0.02, p = 0.89, Table 1) but was generally larger at 12 months of age (F_1, 271_ = 8.8, p = 0.003). No differences between GBH exposed and control females in yolk mass, shell mass, or egg T3 and T4 concentrations were detected (Table 1).

**Table 1.** Quality of the eggs (egg, yolk, and shell mass; thyroid hormones concentrations: T3 = triiodothyronine, T4 = thyroxine) from GBH (glyphosate based herbicide)-exposed and control females. The egg mass was averaged over all eggs (4 and 12 months of exposure). The other parameters were measured after 4 months of exposure.

| **Treatment** | **GBH** | **N**<br>**(GBH)** | **Control** | **N**<br>**(Control)** | **t<sub>df</sub>** | ***p*** |
| --- | --- | --- | --- | --- | --- | --- |
| Egg mass (g) | 10.4 (0.8) | 142 | 10.7 (0.9) | 155 | see text |  |
| Yolk mass (g) | 3.045 (0.296) | 12 | 3.216 (0.254) | 12 | 1.52<sub>22</sub> | 0.14 |
| Shell mass (g) | 1.186 (0.122) | 12 | 1.156 (0.147) | 12 | 0.53<sub>22</sub> | 0.59 |
| T4 (pg/mg) | 6.56 (0.96) | 12 | 6.00 (1.66) | 11 | 1.05<sub>21</sub> | 0.30 |
| T3 (pg/mg) | 5.80(2.18) | 11 | 4.6(1.6) | 11 | 1.45<sub>20</sub> | 0.16 |

Embryo development was normal in 89% of control eggs while 76% of GBH eggs had normally developed embryos. This difference tended to be statistically significant (treatment F_1, 22_ = 3.08, p = 0.09) and the trend was similar at both 4 and 12 months of exposure (treatment*period F_1, 312_ = 0.6, p = 0.43, Figure 1). The un/underdeveloped eggs were distributed across pairs and for none of the pairs were all eggs classified as undeveloped. We detected no major developmental deformities by visual screening of the 10-d embryos and histological samples (see Supplementary Figures 1a, b). Brain mass did not differ between embryos from GBH-exposed and control parents (mean±SD in mg; GBH: 67.1±12.5, control 68.1±15.5; F_1,31_ = 0.04, p = 0.84). Brain oxidative status at 12 months of parental exposure was measured from 19 control and 16 GBH embryos. We measured ca 20% higher lipid damage in the GBH embryos than controls. This difference tended to be statistically significant (F_1, 16.8_ = 3.2, p = 0.088, Table 2), yet there were no differences in the activity of antioxidant enzymes GST, GP, or CAT between the two groups (Table 2).

**Figure 1.**
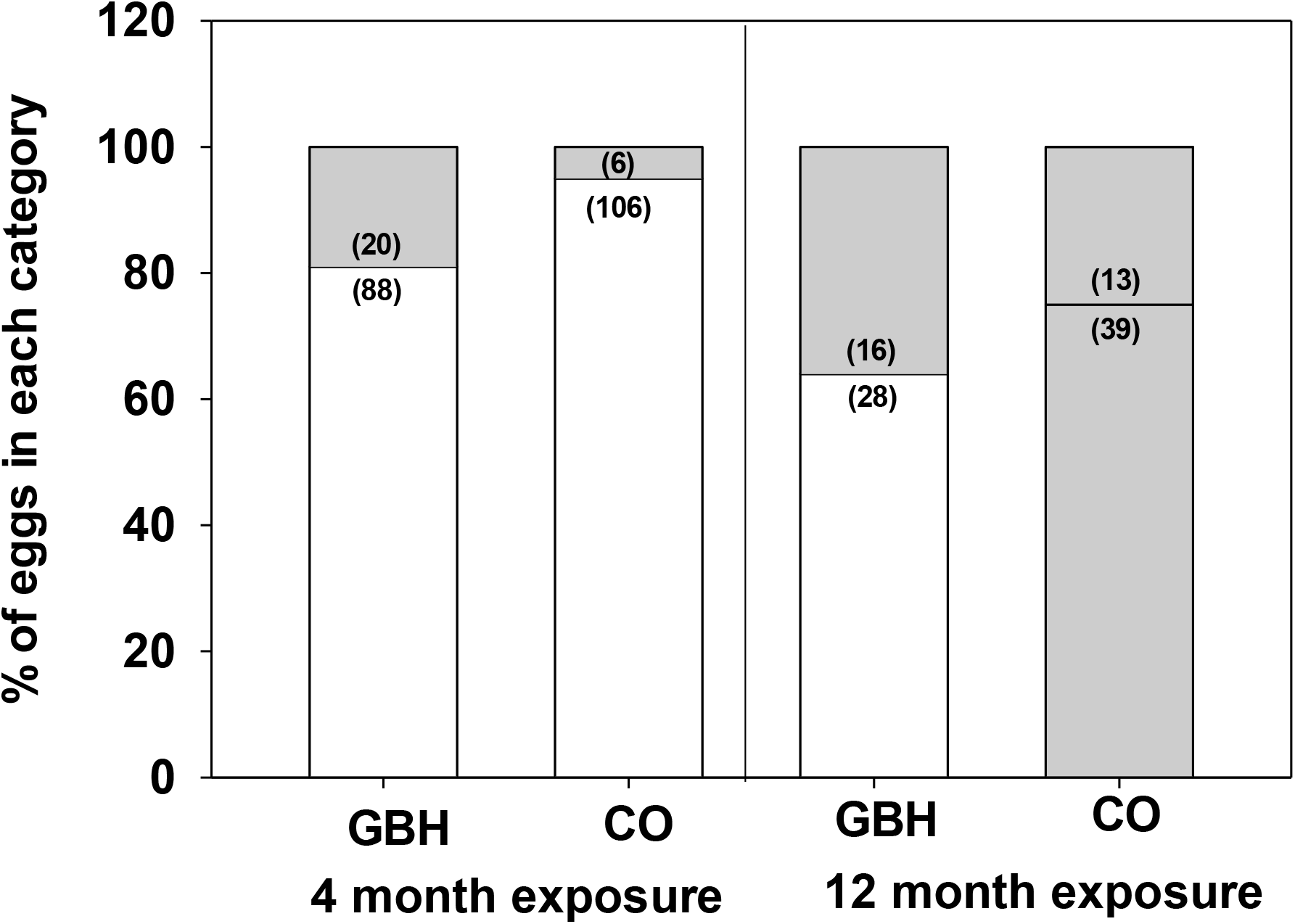
Embryonic status in relation to glyphosate-based herbicide exposure and duration of the exposure. GBH = glyphosate exposed, CO = controls. The percentages refer to percentages of eggs classified in two categories: grey bars = no/little development, white bars = normal development. The bars are drawn separately for GBH and control eggs and after 4 and 12 months of exposure. Sample sizes are indicated in parentheses.

**Table 2.** Average (±SD) of glutathione-S-transferase (GST), glutathione peroxidase (GP), catalase (CAT) activity, and damage to lipids (MDA) in 10-day-old Japanese quail embryos exposed to maternally-derived glyphosate-based herbicide (GBH) or unexposed embryos (control). Associate statistics from linear mixed models (LMMs) are reported below.

| **Treatment** | **GST ($\mu\text{mol/mg}$)** | **GP ($\text{nmol/min/mg}$)** | **CAT ($\mu\text{mol/min/mg}$)** | **MDA ($\mu\text{mol/mg}$)** | **N** |
| --- | --- | --- | --- | --- | --- |
| GBH | 0.0154 (0.004) | 21.73 (4.91) | 6.56 (2.96) | 0.062 (0.016) | 16 |
| Control | 0.0146 (0.003) | 22.08 (4.28) | 6.62 (2.29) | 0.051 (0.018) | 19 |
| LMM | $F_{1, 15.8} = 0.32$ | $F_{1, 33} = 0.05$ | $F_{1, 33} = 0.01$ | $F_{1, 16.8} = 3.2$ | |
| Statistics | $p = 0.57$ | $p = 0.82$ | $p = 0.92$ | $p = 0.088$ | |

## Discussion

Our results indicate that parental exposure to GBHs may lead to negative effects on embryo development and physiology. Because we detected no changes in egg quality (egg, yolk, shell mass, or egg hormone concentration), our results suggest that poorer embryo development and increased brain tissue lipid damage are likely to be explained by the direct effects of GBHs rather than by indirect effects via the altered allocation of resources or hormonal signals to offspring.

The tendency for poorer embryo development in eggs of GBH-exposed parents may be explained by GBH-related effects via either a paternal or maternal route, or both. The eggs with no development visible to the naked eye could have been completely infertile or showing developmental arrest at an early stage. We selected embryonic development as an estimate of reproductive success because it is relevant for the poultry industry and for assessing GBH effects on population growth in wild species. Thus, our methodology could not distinguish between the alternative (paternal versus maternal) underlying mechanisms. Potential infertility may result from GBH-related problems in semen quality or altered reproductive behavior in males as suggested previously in other vertebrates (Cai et al., 2017; Gill et al., 2018; Johansson et al., 2018; Romano et al., 2012). Egg quality was not affected by GBHs and thus the negative effects of GBHs on female folliculogenesis and ovary development, detected in previous studies (Alarcon et al., 2019; Hamdaoui et al., 2018; Schimpf et al., 2017) are unlikely. The fact that non/underdeveloped eggs were scattered across the breeding pairs likely also points to developmental problems of individual embryos mediated via maternally transferred glyphosate to eggs.

Our results support those of previous studies of poultry suggesting that maternal transfer of GBHs to offspring substantially influences offspring via the altering of cell oxidative status. Studies with chicken embryos experimentally exposed to GBHs have shown teratogenic effects on early embryonic development, lower hatchability, and altered serum chemistry (Fathi et al., 2019; Paganelli et al., 2010). Recently, decreased antioxidant enzymes activities (GP, SOD, and CAT) and increased damage to lipids, liver tissue, and kidney tissue post-hatching have been detected (Fathi et al., 2019). In parallel, we observed increased lipid damage in brain tissue, yet antioxidant enzyme activities were not altered. However, it must be noted that the abovementioned experimental studies using *in ovo* injections, researchers applied larger GBH doses compared to those in our experiment (Fathi et al., 2019 ca 10mg/kg, >10x higher than in our study; Paganelli et al., 2010 ca 50mg/kg, i.e. >50x our study), which may explain the different responses. The embryonic brain can be particularly susceptible to ROS because the detoxification system via antioxidants is underdeveloped and polyunsaturated fatty acids are abundant in brain tissue. Indeed, the absence of an effect of GBHs on antioxidants in our experiment may be due to the incapability of the underdeveloped embryonic antioxidant system to respond to GBHs. Oxidative damage to lipids may have serious consequences on brain development (Roy et al., 2016); it is also linked to aging and the onset of many diseases (Simonian & Coyle, 1996). Indeed, GBHs have been shown to have neurotoxic effects on animals such as rodents (Bali et al., 2017; Szekacs & Darvas, 2018).

### Conclusions

In short, our results suggest altered embryonic development and increased embryonic oxidative stress in response to parental GBH exposure in a bird model. Similar results have been found with other vertebrates in previous studies. In a natural ecosystem, transmissive and transgenerational effects may lead to delayed and cascading impacts of agrochemicals; this may potentially explain why some non-target animal populations recover slowly after being exposed to environmental contamination. Thus, recurrent changes in wild populations or in production animals often remain unexplained and may be rarely linked to GBH exposure. While we did not detect any GBH-related changes in maternal allocation, to our knowledge, this is the first long-term study demonstrating transgenerational effects of GBHs with birds. More studies are needed for characterizing GBH-associated changes in maternal allocation and epigenetic programming.

## Acknowledgements

We thank Ida Palmroos, Lyydia Laine, Nicolaus Steinmetz, and Lauri Heikkonen for their help during the data collection, and Tiina Henttinen for the help in histology. The study was funded by the Academy of Finland (grant no. 311077 to MH).

## Data availability statement

The datasets generated and analyzed during this study are available as Supplementary datafiles (1-3).

## Author contributions

SR, MR, MU, and MH designed the study. SR, MU, and MR conducted the data collection. SR conducted the analyses and drafted the first version. All authors contributed to manuscript preparation.

## Supplements

**Supplementary Figure 1.**
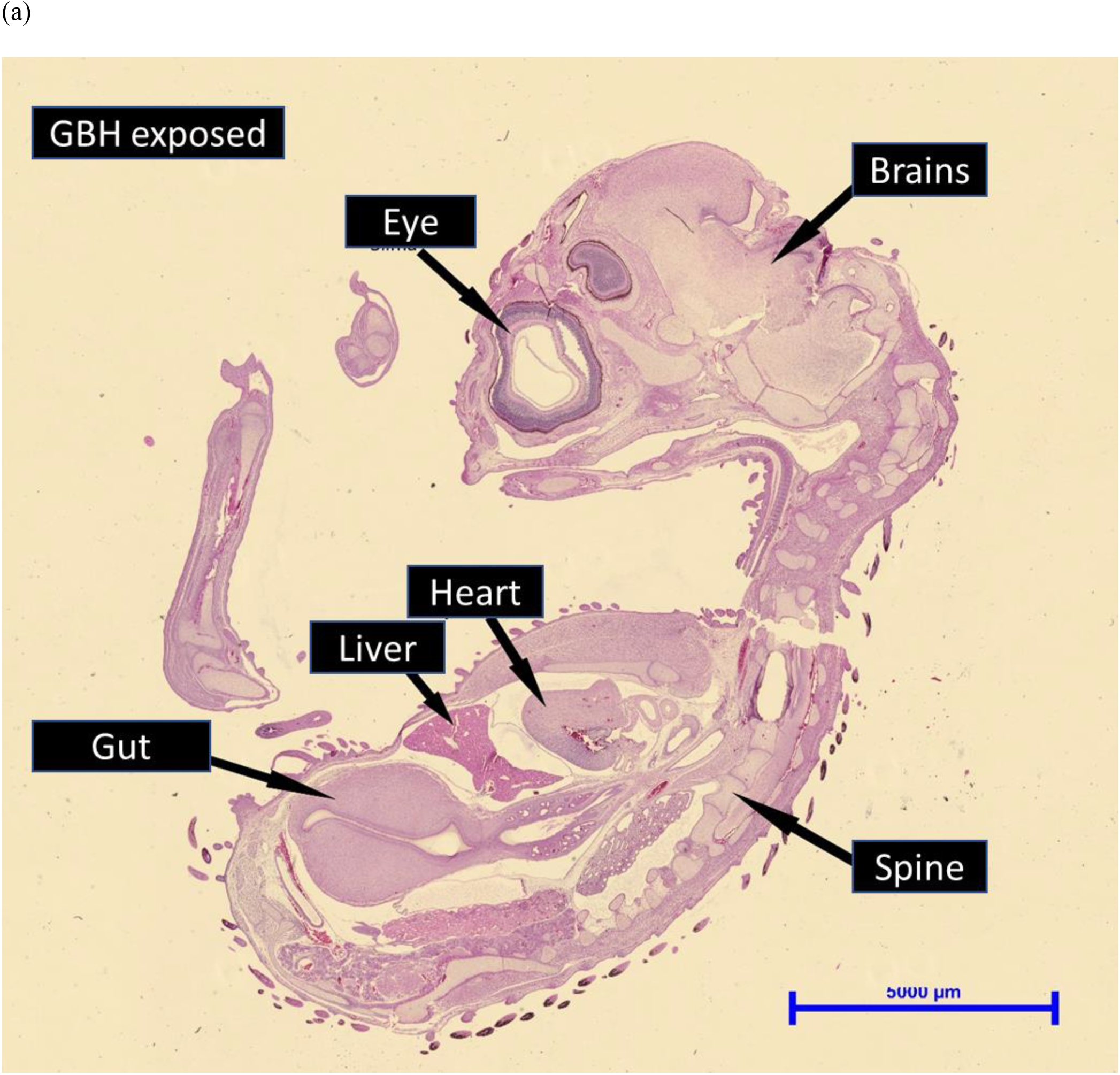
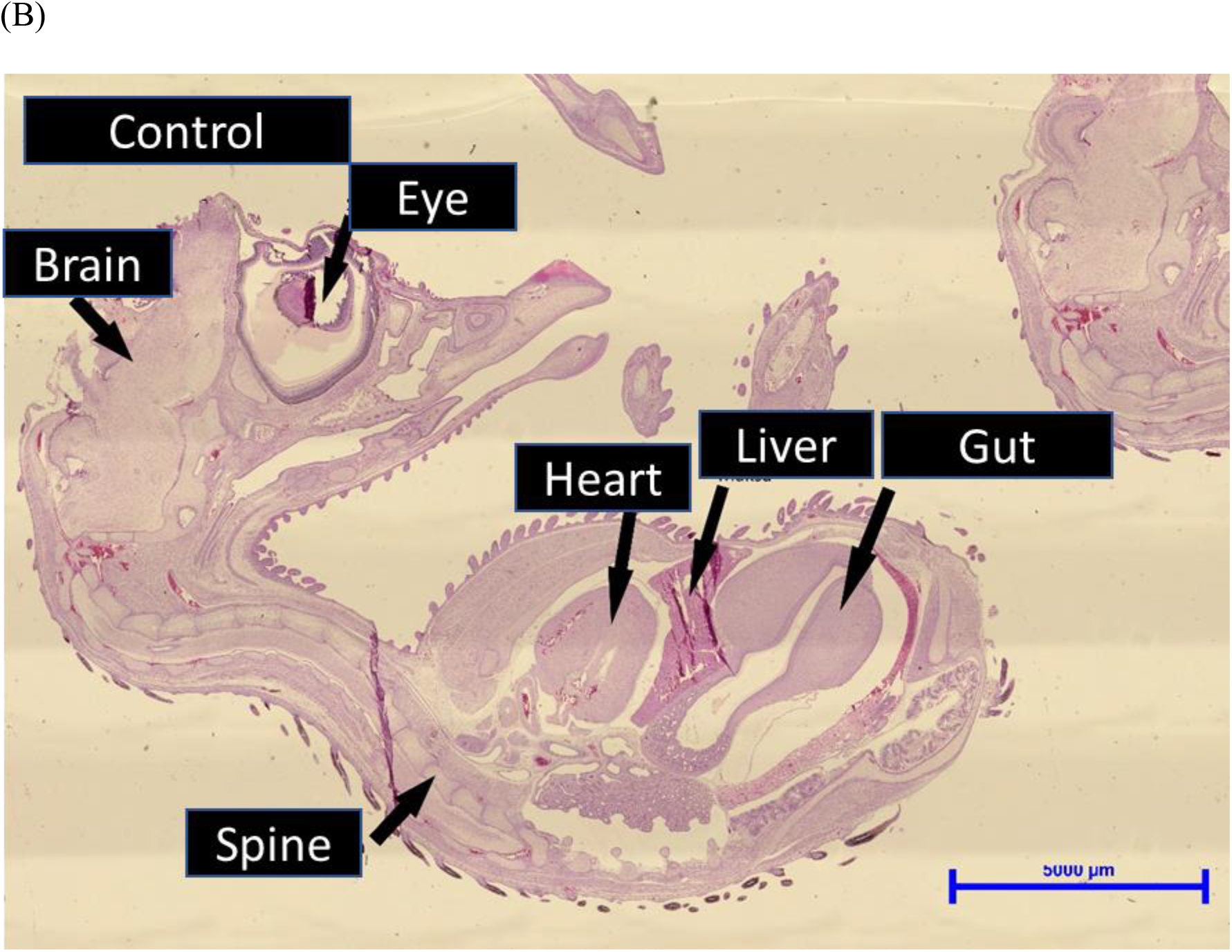
10-day-old Japanese quail embryo (a) from parental generation exposed to glyphosate-based herbicide (GBH) for 10 months; (b) control parents.

